# Identification of circulating protein biomarkers for pancreatic cancer cachexia

**DOI:** 10.1101/359661

**Authors:** Safi Shahda, Ashok Narasimhan, Joshua Kays, Susan M. Perkins, Lijun Cheng, Katheryn N. Hannaford, Daniel E. I. Schloss, Leonidas G. Koniaris, Teresa A. Zimmers

## Abstract

**Background:** Over 80% of patients with pancreatic ductal adenocarcinoma (PDAC) suffer from cachexia, characterized by severe muscle and fat loss. Although various model systems have improved our understanding of cachexia, translating the findings to human cachexia has remained a challenge. In this study, our objectives were to i) identify circulating protein biomarkers using serum for human PDAC cachexia, (ii) identify the ontological functions of the identified biomarkers and (iii) identify new pathways associated with human PDAC cachexia by performing protein co-expression analysis.

**Methods:** Serum from 30 patients with PDAC was collected. Body composition measurements of skeletal muscle index (SMI), skeletal muscle density (SMD), total adipose index (TAI) were obtained from computed tomography scans (CT). Cancer associated weight loss (CAWL), an ordinal classification of history of weight loss and body mass index (BMI) was obtained from medical record. Serum protein profiles and concentrations were generated using SOMAscan, a quantitative aptamer-based assay. Ontological analysis of the proteins correlated with clinical variables (r≥ 0.5 and p<0.05) was performed using DAVID Bioinformatics. Protein co-expression analysis was determined using pairwise Spearman’s correlation.

**Results:** Overall, 111 proteins of 1298 correlated with these clinical measures, 48 proteins for CAWL, 19 for SMI, 14 for SMD, and 30 for TAI. LYVE1, a homolog of CD44 implicated in tumor metastasis, was the top CAWL-associated protein (r= 0.67, p=0.0001). Other proteins such as INHBA, MSTN/GDF11, and PIK3R1 strongly correlated with CAWL. Proteins correlated with cachexia included those associated with proteolysis, acute inflammatory response, as well as B cell and T cell activation. Protein co-expression analysis identified networks such as activation of immune related pathways such as B-cell signaling, Th1 and Th2 pathways, natural killer cell signaling, IL6 signaling, and mitochondrial dysfunction.

**Conclusion:** Taken together, these data both identify immune system molecules and additional secreted factors and pathways not previously associated with PDAC and confirm the activation of previously identified pathways. Identifying altered secreted factors in serum of PDAC patients may assist in developing minimally invasive laboratory tests for clinical cachexia as well as identifying new mediators.

## Introduction

Cancer cachexia is a multifactorial paraneoplastic syndrome characterized by severe loss of muscle and fat leading to overall weight loss [1-3]. Cachexia is commonly seen in patients with advanced cancer and more than 80% of patients with pancreatic ductal adenocarcinoma (PDAC) are affected by this debilitating condition. It has been shown that close to 70% of newly diagnosed cases of pancreatic cancer present with cachexia [4]. Patients with cachexia have decreased survival, increased drug toxicity and decreased physical function, leading to reduced quality of life [5]. Currently, there are no approved therapies for cachexia and it continues to represent an unmet medical need. There has been a surge in developing and studying new models of pancreatic cancer cachexia [6]. Using such model systems, considerable progress has been made in understanding various molecular mechanisms potentially involved in human cancer cachexia [7]. Based upon a myriad of model systems, it is known that cachexia is a result of complex host-tumor interactions, which cannot be simply explained by decreased caloric intake [8, 9]. One of the key drivers of cachexia is systemic inflammation [10, 11]. Nonetheless, understanding the human relevance of these pre-clinical models in human cancer cachexia patients, particularly those suffering with pancreatic cancer, remains poorly defined. To date translating model systems to understand cachexia and develop biomarkers have proven challenging, partly because cachectic patients may be limited in their ability to participate in research trials [12].

Efforts to identify biomarkers for cancer cachexia continue, as it has great diagnostic and therapeutic implications. The use of serum samples to identify biomarkers for human PDAC cachexia should be a possibility in these patients which is minimally invasive. Furthermore, biomarkers should contribute to further understanding of the mechanistic role of serum factors in cachexia. Current practices utilize weight loss as a measurement of cachexia severity; however, readily available Computed Tomography (CT) scans allow us to further quantify the degree of wasting and the compartment being involved (i.e. muscle vs. fat vs. both) [2, 3]. Additionally, it is likely that cachexia is driven by various biological pathways that are disease specific and attempting to group several illnesses would likely confound the results and challenge the ability to draw any meaningful conclusions. Due to the challenges in studying human cancer cachexia, utilizing a minimally invasive procedure to collect tissue (serum) and maximize the utility of available data from clinical records including CT scans, will optimize enrollment to cachexia studies and minimize the burden on patients.

In this study, we sought to identify circulating protein biomarkers for PDAC cachexia. Our objectives were to (i) identify protein biomarkers for cancer cachexia from the serum of patients with PDAC, (ii) identify the ontological functions and canonical pathways of the identified biomarkers and (iii) perform protein co-expression analysis to identify novel pathways associated with cancer cachexia.

## Materials and Methods

### Recruitment of study participants

This was a prospective, observational study that was approved by the Indiana University Institutional Review Board (IRB). The study participants diagnosed with either local or metastatic PDAC were recruited from Indiana University Hospital between the years 2015 and 2017. Written informed consent was obtained from patients for blood and clinical data collections. Patients had to be >18 years of age, provide informed consent, had either confirmed PDAC (advanced group) or at least high suspicion or confirmed PDAC (surgical group). Study procedure including blood samples and clinical data collection were coordinated to meet the standard of care procedures per the treating physician’s discretion. Patients were excluded if they had known HIV or other active malignancies other than PDAC. Serial blood samples were collected from the patients followed for advanced PDAC; and prior to surgery, and during postoperative standard follow up for patients with resectable PDAC. For patients receiving neoadjuvant therapy for resectable PDAC, none received chemotherapy within 3 weeks from the operation. The collected blood was stored in -80C until further use. A total of 30 patients with PDAC, including 19 localized PDAC and 11 with metastatic disease were included in this study. All experiments were performed in accordance with the IRB protocol.

### Assessment of clinical variables for cachexia

Weight loss information over the preceding six months were collected from the electronic medical records, and patients were classified as cachectic if they reported more than 5% weight loss from baseline. Cancer associated weight loss (CAWL), an ordinal classification of history of weight loss and BMI that involves grading of patients from 0 to 4, was calculated for all patients using Martin et al classification [3]. CT scans obtained as part of standard of care follow up and in intervals of every 8-12 weeks were retrieved for body composition analysis utilizing SliceOMatic software; the third lumbar vertebrae were used as a standard landmark to measure the skeletal muscle and total adipose components [2, 3]. Skeletal muscle index (SMI) and total adipose index (TAI) were calculated by normalizing the skeletal muscle and adipose tissue area to their stature (cm^2^/m^2^) [2, 3, 13]. The skeletal muscle density (SMD) was also measured. Sarcopenia status of these patients was calculated based on Martin et al classification [2].

### Protein measurements using SOMAscan

The protein measurements were performed using the serum samples. The SOMAscan proteomic assay is described in more detail elsewhere [14]. In brief, SOMAscan is an aptamer-based technology that utilizes single-stranded DNA aptamers which is modified chemically to enhance the binding to protein epitope with high specificity. Each of the 1,310 proteins measured in serum by the version of the SOMAscan assay performed in this study has its own targeted SOMAmer reagent, which is used as an affinity binding reagent and quantified on a custom Agilent hybridization chip.

Cases associated with early and advanced stage cancer were randomly assigned to plates within each assay run along with a set of calibration and normalization samples. No identifying information was available to the laboratory technicians operating the assay.

Intrarun normalization and interrun calibration were performed according to SOMAscan v3 assay data quality-control procedures as defined in the SomaLogic good laboratory practice quality system. Data from all samples passed quality-control criteria and were fit for analysis.

1310 proteins were assayed with few proteins having more than one probe to bind. The output for every protein from the array is given as relative fluorescence units (RFU), which is proportional to the amount of target protein present in the sample. Hybridization normalization was performed to reduce the technical variation. The data was then median normalized to remove any variation between samples and to account for any variation in the assay. After all the preprocessing steps, quantile normalization was performed across all samples for proteins which will be used for downstream analyses.

### Statistical analyses

The data is represented as mean ± standard deviation, unless otherwise mentioned. Partial Spearman’s correlations adjusted for age and sex were computed to correlate protein expression with CAWL, SMI, SMD and TAI. GraphPad Prism 7 (GraphPad Software, Inc., La Jolla, CA, USA) and R statistical program were used for statistical analyses.

### Functional enrichment analyses for proteins

Proteins correlated with CAWL, SMI, SMD or TAI at a pre-defined cut-off (effect size ≥ 0.5 and p<0.05) were used for functional enrichment analysis, performed using DAVID bioinformatics v7 [15]. Clusters with more than 1.5 enrichment value and p< 0.05 were considered significant.

### Protein co-expression analysis to identify novel pathways associated with cachexia

To identify novel pathways that can potentially be associated with cachexia, in-silico protein co-expression analysis was performed using Spearman correlation. 1,294 proteins were correlated against each other and interactions with ≥ 0.6 (r-value) and p< 0.05 were further considered for interpretation. Pathways of the co-expression network were identified using Ingenuity Pathway Analysis (IPA).

## Results

### Patient demographics/clinical characteristics and Somascan quality control results

The results were reported as mean ± standard deviation (Table 1). In body composition measurements, SMI, SMD and TAI were predominantly reduced when compared against the reference range proposed by Martin et al, suggesting sarcopenic and/or weight loss condition. Percent change in weight was more than 10%, with the metastatic PDAC patients losing more weight than the local PDAC patients (% mean, in six months’ time). As age and sex were shown to significantly alter the expression patterns [16, 17], we have adjusted for age and gender in all our protein correlation analyses. One participant did not have the CAWL grade and two participants did not have CT scans. Therefore, 29 subjects were used for CAWL correlation analysis and 28 subjects for body composition correlation analysis. After passing through the quality control and other pre-processing steps, 16 proteins out of 1,310 did not satisfy the threshold values. In all, 1,294 proteins were subjected to quantile normalization and used for downstream analysis.

**Table 1.** Patient Demographics. Values are indicated as mean ± standard deviation, unless otherwise mentioned. Sarcopenia status and CAWL were assigned using Martin et al. classification.

| Characteristics | PDAC (n=30) |
| --- | --- |
| Age | 67 ± 12 |
| *Gender* |  |
| Male | 16 |
| Female | 14 |
| BMI (kg/m<sup>2</sup>) | 27 ± 6 |
| *CAWL* |  |
| Grade 0 | 3 |
| Grade 1 | 2 |
| Grade 2 | 7 |
| Grade 3 | 8 |
| Grade 4 | 9 |
| *Sarcopenia Status* |  |
| Yes | 20 |
| No | 8 |
| Information not available | 2 |
| Change in weight (% mean) | -10.5 ± 8.5 |
| Skeletal muscle index (cm<sup>2</sup>/m<sup>2</sup>) | 44.6 ± 11.3 |
| Skeletal muscle density (HU) | 27.8 ± 10.1 |
| Total Adipose Index (cm<sup>2</sup>/m<sup>2</sup>) | 146 ± 70.3 |

### Proteins associated with clinical variables of cachexia

As mentioned earlier, proteins with correlation value of ≥ 0.5 and p < 0.05 with clinical variables were considered for interpretation. For identifying biomarkers, the combination of effect size and p-values are considered robust than using p-values alone [18]. At the given cut-off, 48 proteins were correlated with CAWL, 19 with SMI, 14 with SMD, and 30 with TAI. Of the 48 proteins associated with CAWL, 28 proteins were negatively correlated with and 21 proteins were positively correlated. LYVE1, a homolog of CD44 was identified as the top correlated protein with CAWL (r= -0.67, p= 0.0001). Other inflammation related proteins such as C7 (r= 0.65, p=0.0003), C5 (r= -0.55, p= 0.0027), F2 (r=-0.64, p= 0.0003), lymphocyte surface antigen LY9 (r= 0.56, p=0.0022), IFNAR1 (r= 0.51, p= 0.0065) and IL1RL1 (r= 0.50, p= 0.0075) also correlated with CAWL. Proteins that were previously shown to be involved in cachexia such as MSTN (-0.54, p= 0.0040), INHBA (r= 0.50, p= 0.0077) and ALB (r= -0.56, p= 0.0025) were also identified. Interestingly, PH related protein CA10 (r= -0.57, p=0.0021) and stem cell marker NANOG (r= -0.54, p= 0.0038) were also found to be associated with CAWL. The complete list of correlated proteins with CAWL is given in Table 2. The top 30 correlated proteins are represented in Figure 1.

**Figure 1.**
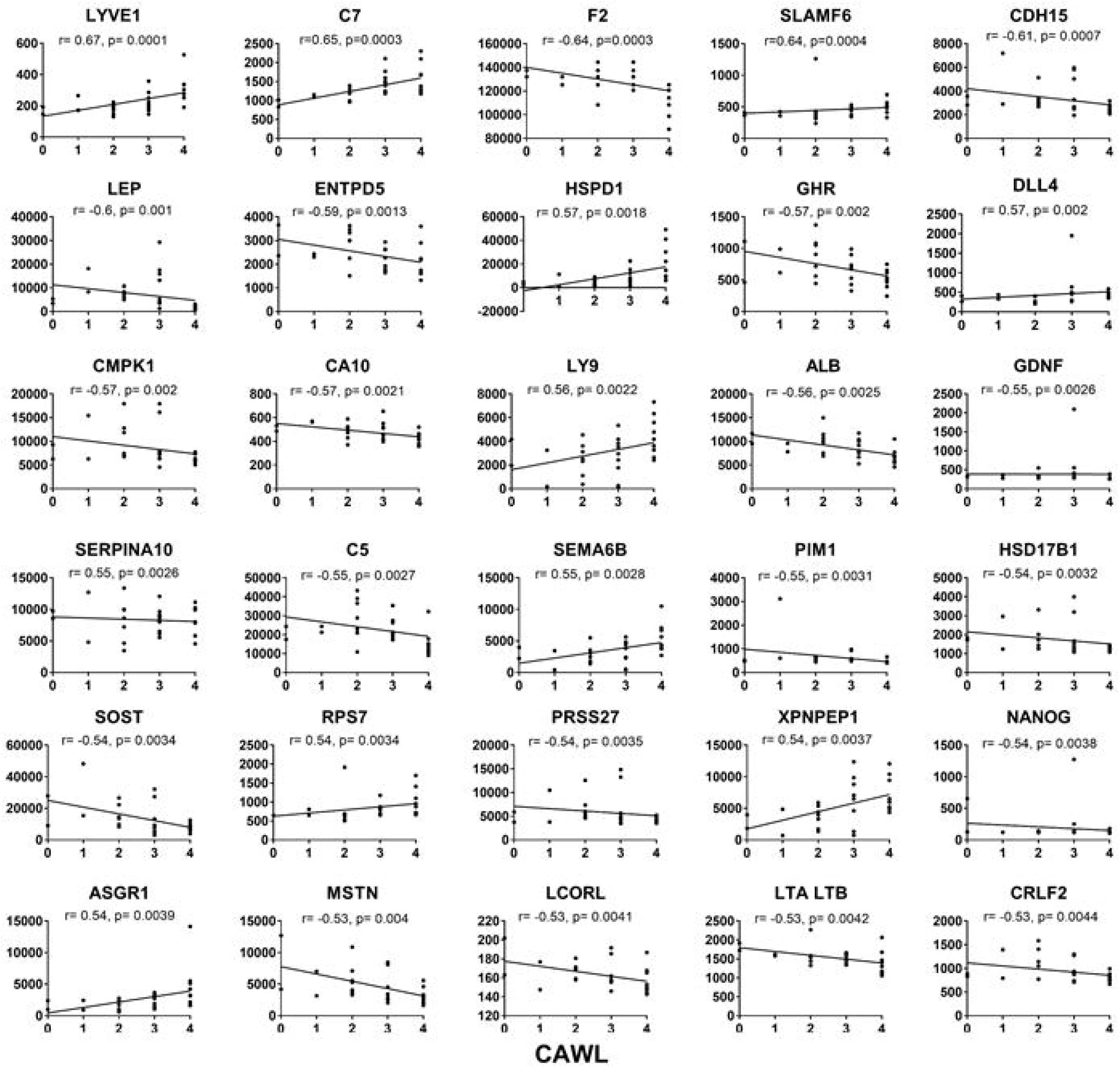
Top 30 proteins correlated with CAWL: The partial spearman correlation r-value adjusted for age and sex and p-values are given for each protein.

**Table 2.**
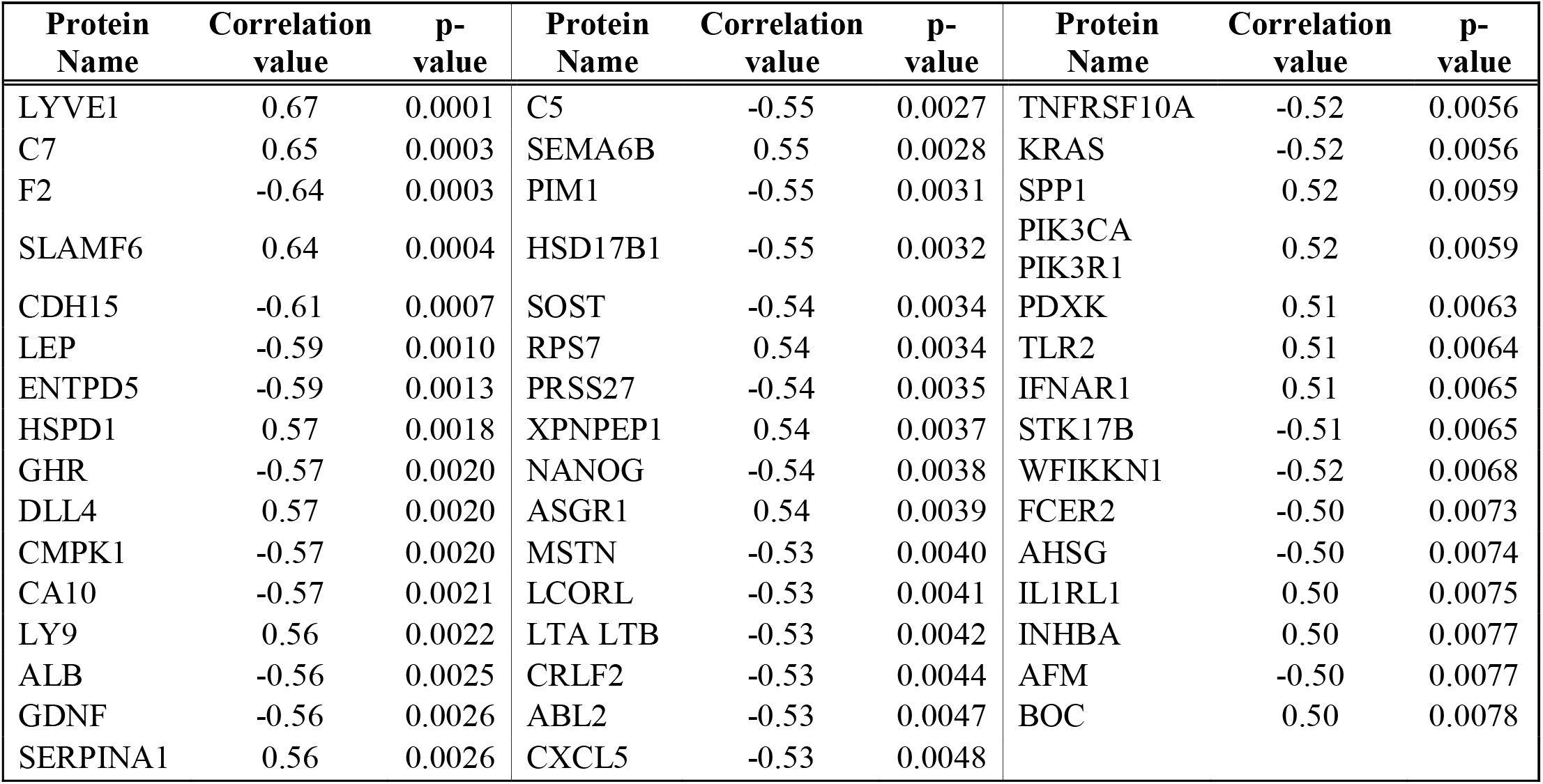
Proteins correlated with CAWL. Partial Spearman Correlation adjusted for age and gender was calculated for CAWL. All proteins which had r ≥ 0.5 and p < 0.05 are reported in this table.

The top correlated protein with SMI was gastrin releasing peptide, GRP (0.70, 0.0001). Other proteins correlated with SMI include acetylation proteins SET (r= 0.56, p= 0.0028) and HDAC8 (r= 0.51, p= 0.0076), inflammatory proteins CFH (r= 0.60, p= 0.0012) and IL1R2 (r= 0.53, p= 0.0054) and calcium binding protein S100A7 (r= 0.62, p= 0.0007). The list of proteins correlated with SMI is presented in Table 3.

**Table 3.** Proteins correlated with SMI. Partial Spearman Correlation adjusted for age and gender was calculated for SMI. All proteins which had r ≥ 0.5 and p < 0.05 are reported in this table.

| **Protein Name** | **Correlation value** | **p-value** | **Protein Name** | **Correlation value** | **p-value** |
| --- | --- | --- | --- | --- | --- |
| GRP | 0.70 | 0.0001 | SET | 0.56 | 0.0028 |
| S100A7 | 0.62 | 0.0007 | LEP | 0.56 | 0.0028 |
| CCL28 | -0.61 | 0.0009 | RSPO3 | -0.56 | 0.0031 |
| CFH | 0.60 | 0.0012 | WFIKKN1 | 0.54 | 0.0041 |
| AFM | 0.59 | 0.0012 | LPO | -0.53 | 0.0050 |
| MB | 0.59 | 0.0015 | IL1R2 | 0.53 | 0.0054 |
| TYMS | 0.57 | 0.0023 | HRG | 0.52 | 0.0066 |
| GHR | 0.57 | 0.0025 | HDAC8 | 0.51 | 0.0076 |
| PIM1 | 0.57 | 0.0026 | MDK | -0.50 | 0.0086 |
| KLK11 | -0.56 | 0.0027 |  |  |  |

FABP3, a fatty acid binding protein involved in fatty acid transport had the highest correlation (r= -0.77, p= 3.7000E-06) with SMD. Many inflammatory related proteins such as CFH (r= -0.58, p= 0.0018), C5 (r= -0.56, p= 0.0031), IFNA7 (r= -0.55, p= 0.0034), IL17B (r= 0.53, p= 0.0051) and FCER2 (r= -0.51, p= 0.0073) were also associated with SMD. The list of correlated protein is reported in Table 4.

**Table 4.** Proteins correlated with SMD Protein Name. Partial Spearman Correlation adjusted for age and gender was calculated for SMD. All proteins which had r ≥ 0.5 and p < 0.05 are reported in this table.

| Protein Name | Correlation value | p-value |
| --- | --- | --- |
| FABP3 | -0.77 | 3.7000E-06 |
| COL18A1 | -0.63 | 0.0006 |
| LRRTM1 | -0.61 | 0.0008 |
| CFH | -0.58 | 0.0018 |
| AKT2 | 0.57 | 0.0025 |
| CFD | -0.57 | 0.0026 |
| C5 | -0.56 | 0.0031 |
| IMPDH1 | 0.56 | 0.0032 |
| IFNA7 | -0.55 | 0.0034 |
| IL17B | 0.53 | 0.0051 |
| LILRB2 | -0.53 | 0.0052 |
| FCER2 | -0.51 | 0.0073 |
| SIGLEC14 | -0.51 | 0.0074 |
| CLEC1B | 0.50 | 0.0086 |

Leptin (LEP, r= 0.85, p= 3.13 E-08), an important molecule in energy homeostasis [19], a mechanism often perturbed in cachexia was the top correlated molecule associated with TAI. SFRP1(r= -0.53, p= 0.006), a protein involved in increased adiposity, APOE (r=0.5, p= 0.009), a member of lipid metabolism and FABP3 (r=0.67, p= 0.0002) were also associated with TAI. Other proteins include C5 (r= 0.69, p= 0.0001), C1S (r= 0.65, p= 0.0003), F9 (r= 0.53, p= 0.006), CCL 28 (r= -0.68, p= 0.0002), PTN (r= -0.52, p= 0.007), to name a few. The list of proteins correlated with TAI is presented in Table 5. As CAWL is a combination of WL% and BMI, many of the proteins that were associated with CAWL were also identified in CT quantified SMI, SMD and TAI.

**Table 5.**
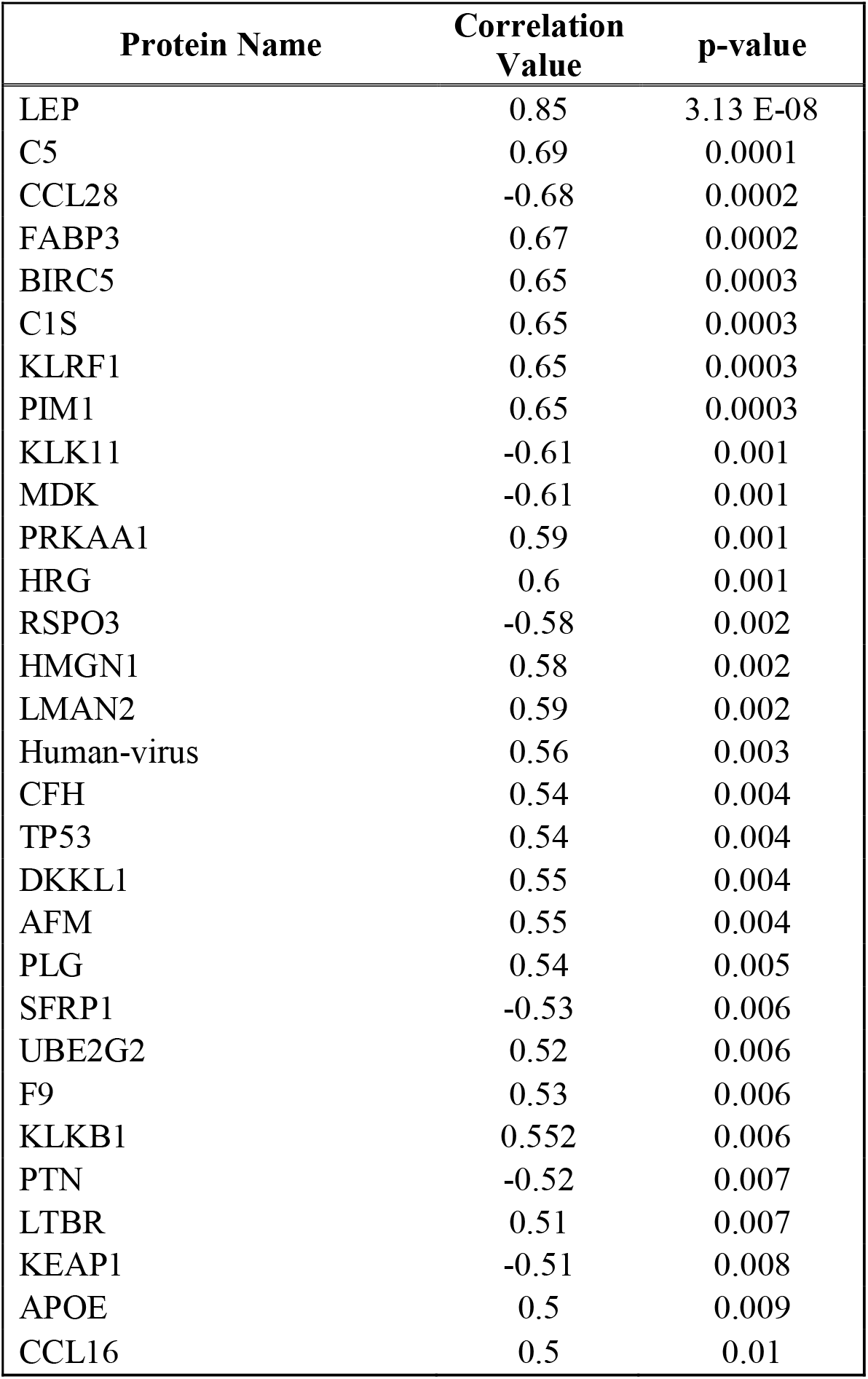
Proteins correlated with TAI. Partial Spearman Correlation adjusted for age and gender was calculated for TAI. All proteins which had r ≥ 0.5 and p < 0.05 are reported in this table.

### Functional enrichment of proteins

The identified proteins that correlated with CAWL, SMI, SMD and TAI were used to identify their functions in PDAC cachexia. Many inflammation related functions such as IL6 production, acute inflammatory response, B-cell activation, T-cell activation, regulation of leukocyte activation, regulation of innate immune response and response to cytokine were identified. This is anticipated as many of the proteins correlated with clinical variables were related to inflammation. We also identified other previously reported functions such as regulation of proteolysis [20], fatty acid transport, response to oxidative stress [21], regulation of JAK-STAT cascade[22], tumor necrosis factor-mediated signaling pathway and MAPK cascade [23] (Figure 2). Similar to our correlation results, we identified certain inflammatory and immune related functions which have not been previously associated with cachexia along with identifying many known pathways and functions in cachexia literature. The complete list of enriched proteins along with their functions is given in Supplementary Table 1.

**Figure 2.**
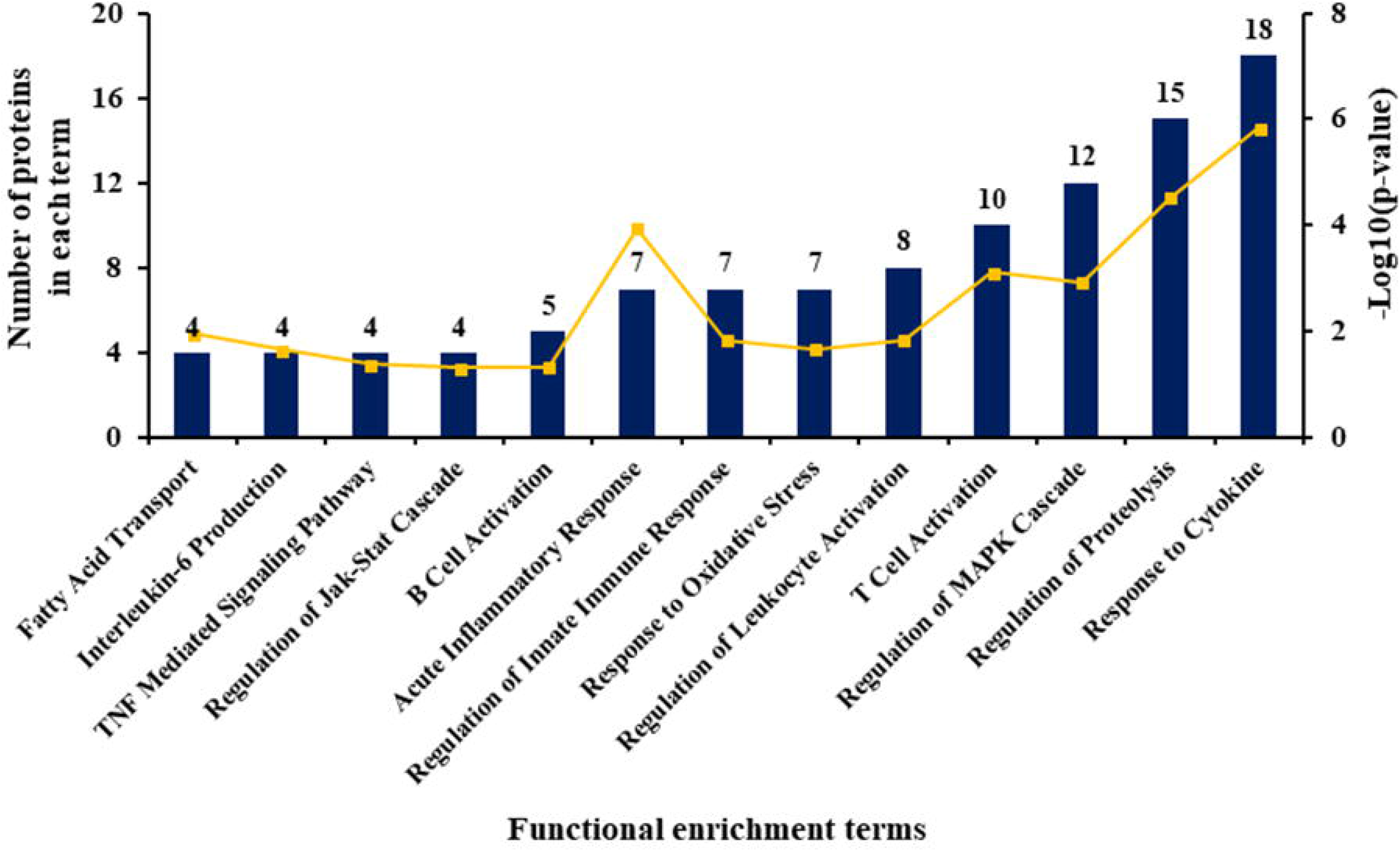
Functional enrichment analysis of proteins correlated with all clinical variables of cancer cachexia: Gene ontology terms enriched for proteins correlated with clinical variables are represented in the X-axis. The number of proteins identified in each pathway is indicated at the top of the bars. Y-axis on the left side indicates the number of proteins and Y-axis on the right side indicates –log10 (p-value) for each of the terms. The yellow line represents the –log10 (p-value).

### Protein co-expression analysis

To identify novel pathways that could potentially be associated with cachexia, we performed a protein co-expression analysis. All 1294 proteins were correlated against each other using Spearman’s correlations. Correlation pairs with r value of > 0.6 and p < 0.05 were considered for the analysis. As CAWL had the highest number of proteins and some proteins associated with SMI, SMD and TAI were also found in CAWL, we took all the CAWL proteins and their correlated pairs to identify the pathways involved in PDAC cachexia. We identified 1,498 correlated pairs for CAWL associated proteins. Non-redundant protein list was imported to IPA and the filters used in IPA were: in prediction category, experimentally validated and highly predicted pathways were selected; in the tissue selections, we used immune cells, skeletal muscle and adipose tissue to identify canonical pathways. Pathways with p<0.05 were considered significant. Representative pathways include B cell receptor signaling, Th1 and Th2 activation pathway, natural killer cell signaling, IL6 signaling, protein ubiquitination pathway, Wnt/β-catenin signaling and mitochondrial dysfunction, PTEN signaling (Figure 3). Other pathways including acute phase response signaling, glucocorticoid receptor signaling, STAT3 pathway, BMP signaling pathway were also identified. Similar to our previous results, many unknown immune system related pathways were identified along with identifying known pathways of cachexia. The complete list of significant canonical pathways is given in Supplementary Table 2.

**Figure 3.**
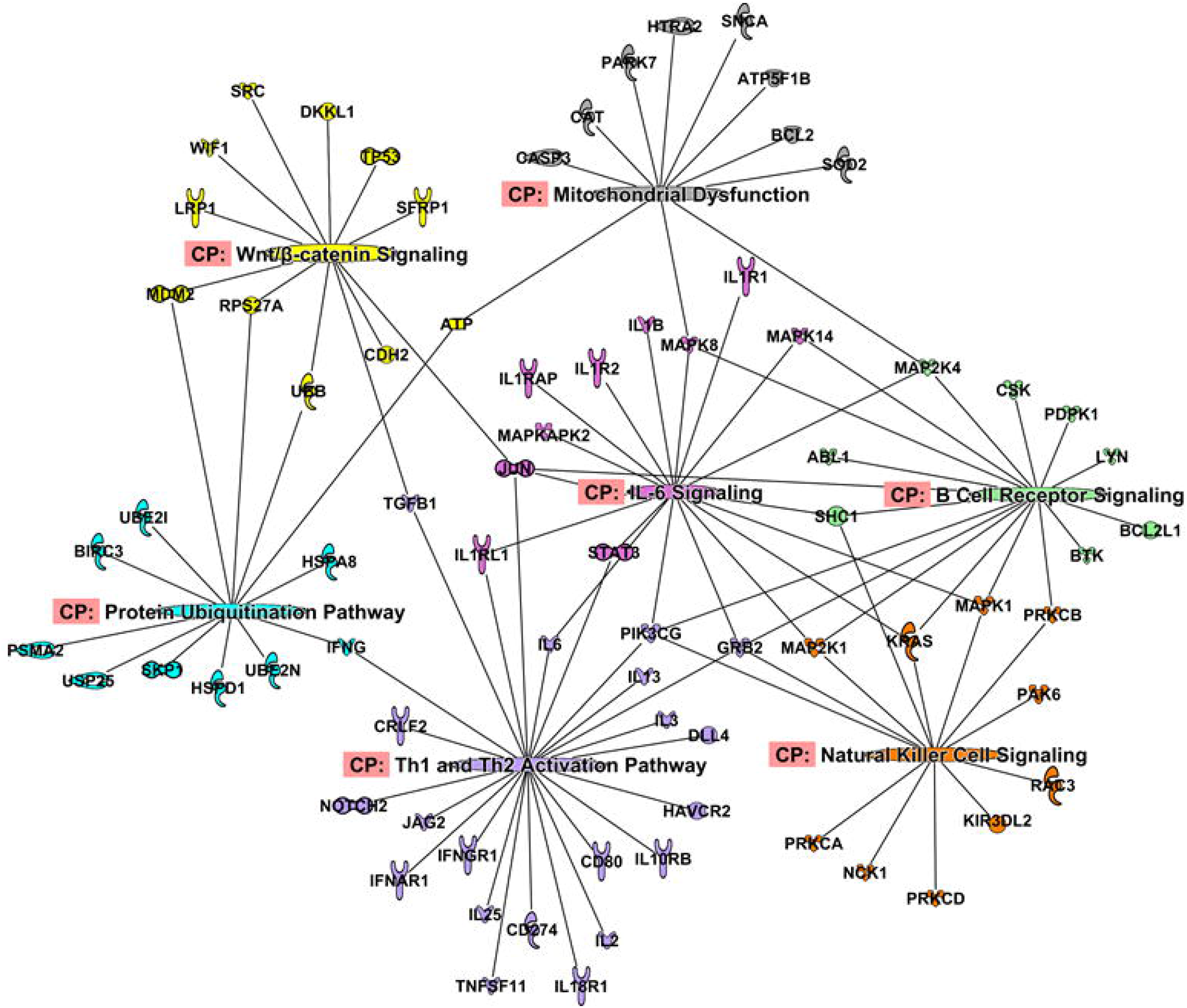
Canonical pathways identified from protein-protein co-expression analysis: Many pathways that were earlier not known in cachexia such as Th1 and Th2 activation pathways, natural killer cell pathways were identified. Some of the well-studied pathways such as IL-6 signaling and protein ubiquitination pathway were also identified.

## Discussion

Using SOMAscan, an aptamer-based assay, we have identified a number of new circulating protein biomarkers associated with cachexia in PDAC patients. This study identifies pathways associated with immune system and inflammation that has not been previously reported in the cachexia literature as makers and potential mediators of PDAC cachexia. One of the strengths of this study is that candidate proteins were identified by an unbiased proteome approach. This allowed independent confirmation of prior studies that used a variety of model systems under one umbrella. Furthermore, it provides a novel ability to examine how many of these pathways may interact with each other in causing and sustaining cachexia. Identifying these pathways using serum samples add to the strength of the study, as it can be used as a noninvasive source to identify candidate pathways from human cachexia and transition towards mechanistically understanding the role of these pathways in cancer cachexia pathophysiology. The availability of comprehensive clinical annotation allowed us to identify potential biomarkers associated with weight loss as well as other cachexia related-body composition measurements. Although weight loss may be an indication of overall body composition changes, not all patients have similar trajectory of muscle and fat loss.

Therefore, identifying biomarkers for the distinct clinical variables of cachexia is a powerful step of building a panel of biomarkers for cachexia diagnosis and therapy.

Our findings suggest that inflammation is a key driver of PDAC cachexia. Functional enrichment and co-expression analysis identified new functions and pathways related to T cells and B cells such as B cell receptor signaling, Th1 and Th2 activation pathways and natural killer cell signaling pathways which we report for the first time in cachexia literature. Other pathways which were previously reported in cachexia such as IL-6 signaling [24], STAT3 signaling [22, 24], protein ubiquitination pathway [20] and mitochondrial dysfunction [25] were also identified.

With cachexia being a manifestation of complex host-tumor interaction leading to muscle wasting and impairing muscle regeneration [8], it is recognized that along with myogenic factors, immune cells have a crucial role in remodeling and regenerating skeletal muscle [11]. The presence of immune cells in healthy skeletal muscle is rare. However, in an injured muscle, the concentration of immune cells increases many fold [26]. T cells were shown to be involved in skeletal muscle regeneration after injury [27]. While Th1 has a proinflammatory effect in recruiting neutrophils and monocytes to the damaged site, Th2 cells promote anti-inflammatory response and myoblast fusion [28]. Therefore, a proper balance between Th1 and Th2 signaling may be crucial for proper muscle regeneration. As the balance in cancer cachexia is shifted more towards proinflammatory than anti-inflammatory effects [29], it remains to be studied if the same mechanism is observed in Th1 and Th2. Alongside, we identified two coagulation proteins F2 and F9 that are correlated with CAWL. Coagulation imbalance leading to excessive thrombosis is one of the complications seen in patients with advanced PDAC. In C26 mouse model of cancer cachexia, hypercoagulation was observed due to partially elevated inflammatory cytokine levels, including interleukin-6 (IL6) [30]. In our study, we found F2 to be negatively correlated with CAWL and F9 to be positively correlated with TAI.

One of the common links between Th1, Th2 pathways and the coagulation proteins F2 and F9 is IL6. IL6 signaling is one of the significant pathways identified in this study and has extensively been studied for its role in PDAC cachexia and as a target for cancer therapy. From our pathway network analysis (Figure 3), IL6 was shown to be involved in Th1 and Th2 activation pathway, and can simultaneously inhibit Th1 polarization and promote Th2 differentiation [31]. Increased levels of IL6 also causes muscle and fat wasting in mice models of cachexia [32]. IL6 and STAT3 were positively correlated with CAWL in our study as well (unpublished observation). These inferences suggest the diverse and critical role of IL6 in cachexia. Therefore, anti-IL6 therapy would be an interesting option to target and see if it can attenuate tumor mass, in turn reducing muscle and fat wasting.

This is the first time that protein biomarkers for myosteatosis are reported. Presence of myosteatosis in patients with cancer severely impacts survival [33]. However, the mechanism through which fat infiltrates in muscle in cancer remains to be elucidated. It remains to be seen if FABP3, a fatty acid transport protein correlated here with SMD may potentially be involved in this process. Additional known inflammatory mediators (ALB, TLR2), and signaling molecules of cachexia (INHBA, MSTN, LEP) were identified in our study. In concordance with what has previously been reported, myostatin and albumin are negatively correlated with CAWL in our study [34, 35].

Although reduced mitochondrial quality has been observed in cachexia preceding the muscle atrophy [25], mechanisms of these events in skeletal muscle needs further investigation. As fatigue is one of the common symptoms in cancer [36], it is not entirely clear whether this is related to muscle wasting leading to fatigue or involves mitochondrial dysfunction as a mechanism of energy wasting. Therefore, understanding these mechanisms may help explore the mitochondrial role in cancer cachexia, and better understand its function in energy homeostasis and muscle dysfunction.

Interestingly, many of the pathways identified from skeletal muscle in earlier studies using experimental models of cachexia have been confirmed in this study from serum samples. This approach could allow us to bridge the gap in understanding the pathophysiology of human PDAC cachexia and accelerate drug development in this devastating condition. Obtaining skeletal muscle biopsies from patients with cancer is an invasive procedure. To understand the disease trajectory of cachexia, collecting serial biopsies at different time points would help better understand the change in muscle microenvironment, however, it is invasive and has to date been difficult to perform. The barriers such as access to muscle samples, the advanced stage of patients with cancer, the focus on cancer therapy and its toxicity have contributed to the slow progress in understanding human cachexia. Using liquid biopsies may prove to be an alternative approach to understand the human pathophysiological changes of cachexia. Further, there are studies to suggest that many of the molecules involved in gene expression, post-transcriptional gene regulatory mechanisms (microRNA and other small non-coding RNA) can be captured using plasma and serum from cancer patients [37, 38]. This remains unexplored for cancer cachexia. Hence, identifying pathways that are enriched in humans, followed by validating in suitable model systems which can capture the heterogeneity of cachexia to an extent could prove to be a powerful strategy for bench to beside approach.

In conclusion, we have identified novel circulating protein biomarkers associated with human PDAC cachexia. We have also identified in the same cohort, previously reported markers of cancer cachexia. We report on inflammatory pathways that were not previously described in cachexia literature. Validation of these findings in an independent PDAC cohort is warranted. It would also be of great interest to explore whether these biomarkers are disease specific by evaluating them in other malignancies associated cachexia.

## Supplementary Information

**Supplementary Table 1. Functional enrichment analysis of proteins correlated with CAWL,SMI, SMD, TAI**

**Supplementary Table 2. Canonical pathways for protein co-expression analysis**

